# Discovery of a phenazine–thiol conjugase from sparse data using genome-informed machine learning

**DOI:** 10.64898/2026.03.05.709892

**Authors:** Xiaoyu Shan, Inês B. Trindade, Nathaniel R. Glasser, Korbinian O. Thalhammer, Matthew Scurria, Ariane Mora, Stuart J. Conway, Dianne K. Newman

**Author notes:** Corresponding author: Dianne K. Newman. These authors contributed equally to this work (Xiaoyu Shan, Inês B. Trindade). **Author Contributions:** X.S, I.B.T and D.K.N designed research. X.S performed genomic data analysis and developed the machine learning framework with significant assistance of A.M. I.B.T performed protein purification and biochemical characterization with assistance of X.S. N.R.G, K.O.T, I.B.T and X.S performed mass spectrometry analysis. M.S and S.J.C contributed to the chemical reaction scheme. X.S and I.B.T performed genetic experiments and phenotypic experiments. X.S. I.B.T and D.K.N wrote the paper with input from all the authors. **Competing Interest Statement:** The authors declare no competing interests.

## Abstract

Machine learning has enabled powerful biological discoveries using models trained on large datasets. However, for many important biological questions, such as identifying enzymes that transform understudied substrates, sparsity of training data is often a major bottleneck. Here, using phenazine natural products as a case study, we show that integrating genome-informed data augmentation with contrastive learning in protein language space enables identification of phenazine-interacting proteins starting from only 14 known phenazine modifying sequences. Applying this framework led to the discovery of PTC (Phenazine-Thiol Conjugase), the first enzyme known to catalyze phenazine thioconjugation, a phenazine modification reaction long observed but previously presumed to occur only through non-enzymatic chemistry. In silico simulation and experimental measurements demonstrate that PTC binds to both phenazine and glutathione as substrates. Recombinant expression and biochemical characterization reveal that PTC promotes glutathione-dependent modification of phenazines, yielding distinct reaction outcomes that depend on substrate identity. Although thiol-conjugated phenazine products exhibit reduced toxicity to bacterial cells, deletion of the gene encoding PTC does not confer a strong fitness disadvantage, illustrating how direct learning of sequences can uncover relevant enzymes that might evade phenotype-based genetic screens. Together, these results demonstrate that coupling comparative genomics with protein machine learning can convert “small data” typically outside the scope of machine learning into actionable predictive power, thereby facilitating enzyme discovery.

**Significance:** Machine learning excels when large, well-labeled datasets are available, yet many biologically important problems lack sufficient experimental data to support such approaches to discovery. This limitation is particularly acute for identifying enzymes acting on rare or understudied substrates. Here, we show that genomic organization can be leveraged as an additional source of biological information to address data sparsity. Starting with only 14 enzymes experimentally shown to modify phenazines, we developed a model identifying phenazine-interacting enzymes by integrating genome-informed data augmentation with protein machine learning. Guided by the model, we discovered the first enzyme known to catalyze thioconjugation modifications of phenazines, demonstrating a simple yet powerful strategy for extracting predictive insight from sparse biological knowledge.

## Introduction

Machine learning has revolutionized biological discovery by extracting predictive power from sufficiently large training data (1, 2). The remarkable success of protein structure prediction, for example, rests on decades of community effort that produced extensive, curated structural datasets (3). In contrast, however, many biologically important questions remain largely inaccessible to machine learning because relevant training data are extremely sparse (4, 5). A prominent example among those is the identification of enzymes that act on specific, understudied substrates, for which only a handful of experimentally characterized sequences exist. In such cases, generating training data through large-scale experimental screening is often slow, expensive, and only feasible when an easily measurable phenotype is available.

Although enzyme function is ultimately encoded in amino acid sequence, in bacterial genomes, genomic organization is itself highly informative (6). Bacterial genomes are structured by evolutionary pressures that favor the co-evolution and co-localization of functionally related genes, giving rise to conserved organizational patterns over evolutionary timescales (7). Ignoring this genomic context discards a substantial source of functional information. A classic example is provided by the catalytic subunits of respiratory nitrate reductase (NarG), which is homologous to nitrite oxidoreductase (NxrA), catalyzing the same reaction but in the opposite direction. They are often difficult to distinguish based on amino acid sequence alone, whereas their genomic contexts (i.e., *narGHI* vs. *nxrABC* operons) easily resolve their functions. Similar principles apply broadly in natural product biosynthesis, where enzymes that modify a shared chemical scaffold are frequently co-localized with the genes encoding the core biosynthetic machinery, together forming biosynthetic gene clusters (8, 9). When such clusters are distributed across diverse phylogenetic lineages, the associated modifying enzymes can diverge extensively in sequence while retaining conserved substrate specificity. The resulting natural exploration of sequence space provides a rich training signal enabling machine learning in the protein language space to extract transferable latent sequence features. When executed systematically, such strategy of combining comparative genomics with protein machine learning has the potential to bring many traditionally data-limited biological questions within the scope of machine-learning-based discovery.

Phenazines provide a representative example of a biologically important yet data-sparse family of microbial natural products (10). Built on a conserved heterocyclic three-ring scaffold, phenazines are produced by diverse bacteria and can be chemically modified into a wide range of derivatives that fulfill distinct roles (11). For instance, pyocyanin (1-hydroxy-5-methylphenazine) is a major virulence factor of the human pathogen *Pseudomonas aeruginosa* (12), whereas PCN (phenazine-1-carboxamide) has been investigated as a biocontrol agent for suppressing fungal pathogens in agriculture (13). Several other phenazine derivatives have attracted substantial interest in biomedical contexts for exhibiting strong antibiotic and antitumor activity (14). Despite their broad clinical and ecological relevance, knowledge of phenazine-modifying enzymes remains limited, with fewer than 20 unique modification enzymes reported with experimental evidence (15). The small size of this training dataset has hindered conventional machine-learning efforts to identify the shared signatures from known phenazine-interacting protein sequences, motivating the development of new strategies enabling prediction directly from small data.

Here, using phenazines as a case study, we demonstrate that genome-informed data augmentation combined with a simple contrastive learning framework enables zero-shot discovery of previously uncharacterized phenazine-modifying enzymes. Starting from 14 known positive sequences, this approach led to the identification of the first enzyme catalyzing phenazine thioconjugation, a detoxification reaction long presumed to occur only non-enzymatically. Overall, the strategy of integrating genome-informed data augmentation and contrastive protein representation promises to unlock hidden enzymatic chemistry encoded in genomes by making specific data-limited questions accessible to machine learning. We thereby term this strategy ML-CITO (<u>M</u>achine <u>L</u>earning for genomic <u>C</u>ontext-Informed <u>T</u>ransferable disc<u>O</u>very).

## Results

### Genomics-informed machine learning enables zero-shot predictions from small data

To enable machine-learning discovery of previously uncharacterized phenazine-modifying enzymes (Figure 1), we first addressed the challenge of limited training data by leveraging comparative genomics. We curated a set of 14 unique (i.e., no homology between them) phenazine-modifying enzymes that were identified experimentally (15), most of which were from phenazine-producing bacteria (Table S1). These phenazine producers synthesize the phenazine core scaffold using a conserved biosynthetic pathway (e.g., core biosynthesis genes *phzABCDEFG* in *Pseudomonas*, Figure1B); the core phenazine structure is then biochemically modified into derivatives. The enzymes catalyzing each step of modification (e.g., adding a methyl group by PhzM or adding an oxygen atom by PhzS, Figure 1B) are often encoded next to the core biosynthetic genes, enabling their identification via conserved genomic context across diverse phenazine producers (16).

**Figure 1.**
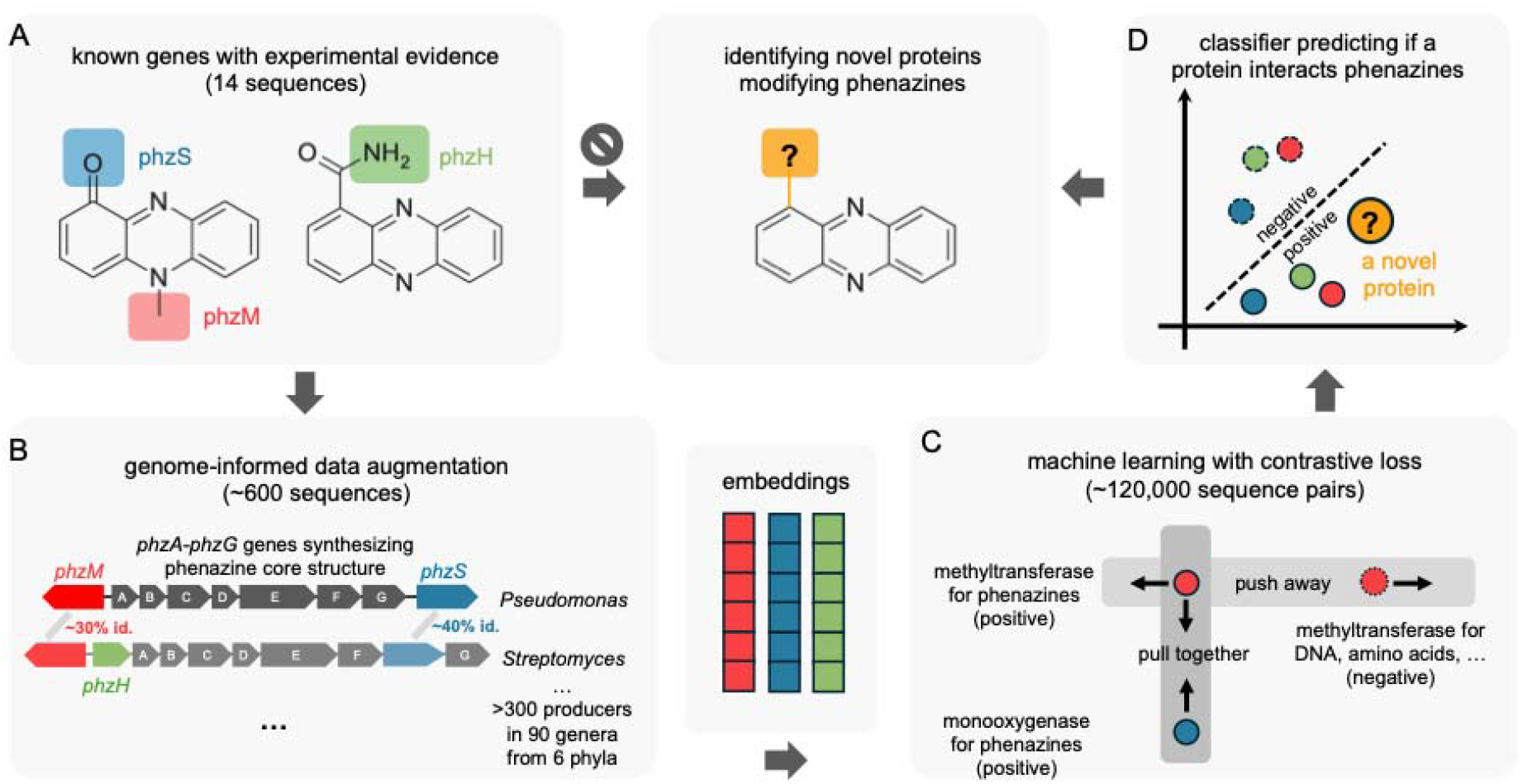
Schematic illustration of ML-CITO framework for phenazine-modifying enzyme discovery. (A) Examples of known phenazine-modifying enzymes with experimental evidence, such as the methyltransferase PhzM, monooxygenase PhzS, and amidotransferase PhzH. (B) Conserved genomic organization of phenazine biosynthetic gene clusters is leveraged to expand 14 unique seed enzymes into an augmented training set of ∼600 homologous sequences of 14 homologous groups. There is substantial sequence variation within each homologous group because of the broad taxonomic distribution of phenazine biosynthesis. (C) Protein sequences are embedded using a pretrained protein language model and passed to a neural network. The model is trained using contrastive learning such that enzymes that act on phenazines but belong to different functional categories are pulled together in latent space, while enzymes from the same functional category that act on phenazines versus other well-studied substrates are pushed apart. (D) The transformed latent embeddings are subsequently classified using a linear support vector machine to distinguish phenazine-interacting enzymes from non-interacting ones. The model enables the prediction of phenazine interaction for any new protein sequence based solely on amino-acid sequence information.

To expand the sparse training dataset of phenazine-modifying enzymes, we surveyed ∼200,000 non-redundant genomes across the Bacteria domain (17). We identified 317 candidate phenazine producers containing the core biosynthetic gene cluster based on both sequence and synteny (Method, Dataset S1). These producers span 90 different genera across 6 bacterial phyla (i.e., Actinomycetota, Chloroflexota, Cyanobacteriota, Myxococcota, Pseudomonadota, Verrucomicrobiota), reflecting substantial taxonomic breadth and evolutionary diversity (Dataset S1). By searching for homologs of the known phenazine-modifying enzymes that also co-localize with core biosynthetic genes, we expanded the initial set of 14 enzymes to ∼600 hypothesized proteins with substantial sequence variation (Methods, Dataset S2). For example, *phzM* from *Pseudomonas aeruginosa* is a biochemically validated phenazine methyltransferase located immediately adjacent to *phzA-G* (Figure 1B). Homologs of *phzM* sharing low sequence identity in distantly related taxa (e.g., 30.3% sequence identity in *Flexivirga sp026162425*, 42.1% sequence identity in *Streptomyces sp018070345*) can nevertheless be confidently assigned as positives based on highly conserved genomic context (Figure 1B). This context-based annotation is further supported by prior experimental studies validating the functions of several such homologs located near the core cluster (18–22). The ∼600 positive sequences augmented from 14 non-homologous seed enzymes span 6 broad functional categories, including amidotransferases, methyltransferases, monooxygenases, prenyltransferases, AMP-binding enzymes, and dehydrogenases (Table S1). To balance the positive sequences for each functional category, we assembled a set of ∼600 negative sequences that are unlikely to modify phenazines but belong to the same functional categories as the positives (e.g., well-studied DNA-modifying methyltransferases vs. phenazine-modifying methyltransferases, Table S2 & S3, Dataset S2). Importantly, negative sequences were also sampled from the same set of phenazine-producing genomes from which the positive sequences derived, minimizing confounding phylogenetic signals in the protein sequences that could otherwise be exploited by the classifier.

With a balanced collection of positive and negative sequences spanning the same enzyme families in hand, we then aimed to distinguish phenazine-modifying enzymes from functionally related but non-interacting proteins using machine learning. To that end, we first embed the protein sequences using a pretrained protein language model (23). To capture substrate-centric signatures of phenazine interactions to transcend the learnt functional categories, we trained a neural network to pull together embeddings of positive pairs drawn from different functional categories (e.g., a phenazine-modifying methyltransferase and a phenazine-modifying monooxygenase, Figure 1C). In the meantime, the model also pushes apart embeddings of positive–negative pairs drawn from the same functional category (e.g., a phenazine-modifying methyltransferase versus a DNA-modifying methyltransferase, Figure 1C). This approach, known as contrastive learning (24), has been widely used as a data-efficient machine-learning method for classification tasks such as enzyme function prediction (25), catalytic residue prediction (26) and enzyme-drug interactions (27). A linear support vector machine was trained on the contrastively transformed embeddings to classify sequences as phenazine-interacting or not, enabling prediction for any new given protein with only its amino acid sequence (Figure 1D).

To assess the predictive capability of the model, we performed three complementary and increasingly challenging evaluations. First, because positive and negative sequences were curated within the same 6 broad enzyme families, we performed leave-one-category-out cross-validation to test zero-shot transfer to entirely unseen functional categories. In each of six tests, all positives and negatives from one category were withheld as test data, and the remaining five categories were used for training. Across folds, the model generalized across categories with performance varying by the held-out family (balanced accuracy = 0.75 ± 0.15; Area Under the Precision-Recall Curve AUPR = 0.81 ± 0.17; mean ± SD across six tests; Figure 2A). We report AUPR, which unlike balanced accuracy provides a threshold-independent assessment of classifier performance and has been widely adopted in enzyme discovery studies (28, 29). Second, we evaluated a more realistic out-of-distribution setting using 10 additional enzymes reported to interact with phenazines yet falling outside the modification families represented in training (e.g., phenazine-degrading dioxygenases and demethylases, and putative phenazine-binding chaperones; no homology with the trained modification sequences; Table S4). These enzymes are limited in augmentation guided by genomic context (7/10 from non-producer genomes) and therefore serve as an orthogonal test of whether the model has learned phenazine-relevant sequence features rather than producer- or cluster-associated artifacts. The classifier correctly identified 8/10 of these proteins as positive, while labeling 9/10 randomly chosen metabolic enzymes from unrelated families as negative (Table S4), indicating generalization capability without reliance on biosynthetic gene cluster context.

**Figure 2.**
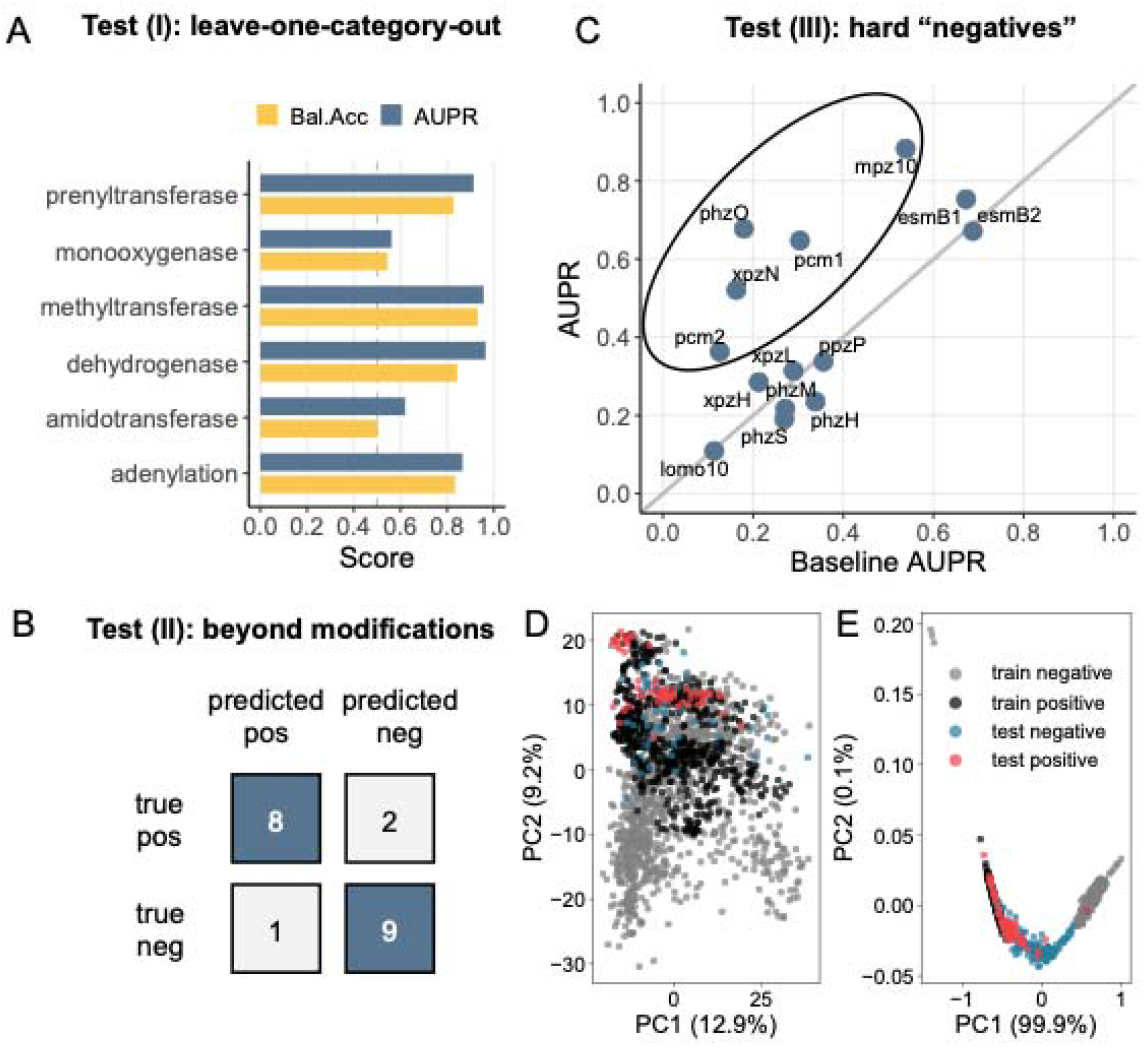
ML-CITO performance evaluation. (A) Leave-one-category-out cross-validation across six broad functional categories (amidotransferases, methyltransferases, monooxygenases, prenyltransferases, AMP-binding enzymes, and dehydrogenases), in which one entire category of positive and negative sequences is withheld for testing while the remaining five are used for training. Bal. Acc, balanced accuracy; AUPR, area under precision-recall curve. (B) Evaluation of ten additional phenazine-interacting enzymes outside the six training functional categories and largely independent of phenazine biosynthetic gene clusters (e.g., phenazine-degrading from non-producer microorganisms), alongside ten well-characterized metabolic enzymes unrelated to phenazines serving as negatives. (C) A deliberately adversarial evaluation in which positive sequences and “hard negative” sequences are classified by ML-CITO: these sequences are homologs to each other, only differing in their genomic distance from the phenazine biosynthetic gene cluster, yet neither group shares homology with any training sequence (see Methods). Despite this challenging setup, five withheld groups show substantial gains in AUPR relative to the baseline null expectation. (D) Embeddings of sequences projected onto a principal component analysis (PCA) ordination plot before training. All homologs of *pcm1* in (C) were left out so they could later serve as the test set. Before training, positive and negative sequences are hard to distinguish. € Transformed embeddings of sequences projected onto a PCA ordination plot after training with contrastive loss, which separates positive (black) from negative (gray) sequences. Applying the trained model to the unseen *pcm1* homologs, positive homologs (red) are also well separated from the negative homologs (blue).

Finally, we stress-tested the model on “hard negatives” by subjecting it to a challenging evaluation. If the model we trained indeed captures sequence signatures specific to phenazine interactions, it should not only discriminate phenazine modifiers within the same broad functional categories but also identify between closely related homologs. To test whether this was the case, we withheld from training all homologs of one of the 14 seed modification enzymes that are located within 10 genes of the core biosynthesis genes. These withheld homologs served as positive test examples, whereas homologs of the same enzyme located more than 50 genes away from the core cluster were designated as “hard negatives” (i.e. unlikely to modify phenazine, see Figure S1 for schematics). We anticipated that our “hard negative” set would comprise not only true negatives but also true positives because some modification genes can be decoupled from the core genes (e.g., *phzH* in *P. aeruginosa*). In addition to the potential for positive contamination of the “hard negatives”, this task was also intrinsically more challenging than the previous category-level tests because the model had to discriminate between positive and “negative” sequences that are homologous to each other yet share no homology with any training sequences. As expected given this challenging setup, the AUPR was near baseline for many groups, reflecting the null expectation; nevertheless, five groups (*mpz10, pcm1, xpzN, phzO, pcm2*) showed substantial improvement over the baseline expectation (ΔAUPR = +0.23 ∼ +0.49; Figure 2C). The increased AUPR indicates that the model ranks the likely phenazine modifiers higher than their homologs that are less likely, suggesting that the model may capture intrinsic sequence features associated with phenazine interactions transcending training sequences. For example, the positive and hard negative sequences homologous to phenazine acyl-CoA dehydrogenase *pcm1* can be separated from each other by the trained model, which has never seen any *pcm1* homologs during its training (see Figure 2D-E, where pre-trained and post-trained embeddings are projected onto a PCA plot with *pcm1* positive and hard negative homologs held out as the test set).

Collectively, the evaluations above show that a simple model, trained from only 14 seed enzymes and augmented through genomic context, can identify phenazine-modifying enzymes across diverse evolutionary, functional, and genomic contexts.

### Identifying the first phenazine thiol conjugase guided by the model

We next applied the trained model to identify previously unknown phenazine-modifying enzymes in bacterial genomes. We selected *Pantoea agglomerans* as a source organism because it is a widespread soil bacterium able to make and transform diverse antibiotics, yet it is underexplored relative to dominant soil taxa such as *Pseudomonas* and *Streptomyces (30, 31)*. Our laboratory strain, *P. agglomerans* W2I1 (NCBI assembly GCF_030815835.1), does not encode phenazine biosynthesis genes; however, it was co-isolated from the wheat rhizosphere together with multiple phenazine-producing bacteria (32), making it likely to harbor enzymes that interact with exogenous phenazines. When all proteins encoded in this genome were scored by the model, 83.5% were consistently assigned negative labels (Dataset S3), as expected for enzymes unrelated to phenazine metabolism. The remaining proteins were assigned with positive scores ranging from 0.001 to 1, and we therefore ranked them by decreasing predicted score (Dataset S3). Among the top ten candidates we noticed a 148-amino-acid uncharacterized protein (UniProt entry ID A0A379LSW4) annotated as a “putative lactoylglutathione lyase” for its weak similarity (27.2% identity) to *Escherichia coli* GloA, a member of the glyoxalase I family that uses glutathione as a substrate. Although this protein is not homologous to any known phenazine modification enzymes, such annotation suggested a possible chemical link to phenazine transformation, as thiol compounds like glutathione often mediate detoxification via thiol adduction (33). Conjugates between phenazines and thiols (e.g., pyocyanin–glutathione adducts) have long been observed, yet these reactions have been assumed to only occur non-enzymatically (34, 35). This inference was made in part because in previous studies where thioconjugated phenazines were detected, no glutathione-related enzymes near phenazine biosynthetic gene clusters were identified (34). Accordingly, we reasoned that if this enzyme were capable of catalyzing phenazine thioconjugation, it would represent the first such enzyme described, motivating us to prioritize its study. Notably, the identification of this enzyme by our model from a genome lacking phenazine biosynthesis genes indicates that the model could be capturing deeper sequence features associated with phenazine interactions rather than relying on biosynthetic gene cluster proximity alone.

With this hypothesis in mind, we explored candidate substrates and cofactors through structural modelling. AlphaFill analysis (36) revealed strong structural similarity to experimentally characterized glyoxalase I enzymes from diverse organisms, including human, murine, and bacterial homologs (PDB IDs: 1QIP, 4MTQ, 1FRO, and 1BH5). In these structures, glutathione and divalent metal ions such as Zn^2^□ or Ni^2^□ are positioned as substrates and cofactors within a conserved active-site pocket. To further examine substrate binding, we generated dimeric structural models of our protein using Chai-1 (37) in the presence of various divalent metals (e.g., Zn^2^□, Ni^2^□) and phenazines (e.g. pyocyanin (PYO), 1-hydroxyphenazine (1-OH-PHZ)). In all cases, the metals consistently occupied the same conserved binding site coordinated by two histidine residues and one glutamate residue (His30, His97, and Glu147), consistent with the canonical His□–Glu motif characteristic of the Vicinal Oxygen Chelate (VOC) superfamily to which glyoxalase I enzymes belong (38, 39). Docking simulations with pyocyanin positioned the phenazine ring near both the glutathione-binding pocket and the metal center, supporting catalytic activity (Figure 3A). To test this hypothesis, the enzyme was recombinantly expressed and purified to >99% homogeneity using immobilized metal affinity chromatography with a stepwise gradient (Figure 3B). Fluorescence quenching experiments confirmed binding of reduced glutathione, yielding a dissociation constant (K_d_) of 9.4 ± 1.8 μM (Figure 3C, Figure S2). STD-NMR further confirmed binding of pyocyanin, with the oxygen-containing ring showing the highest saturation transfer, consistent with its role as the primary epitope (Figure 3D-F).

**Figure 3.**
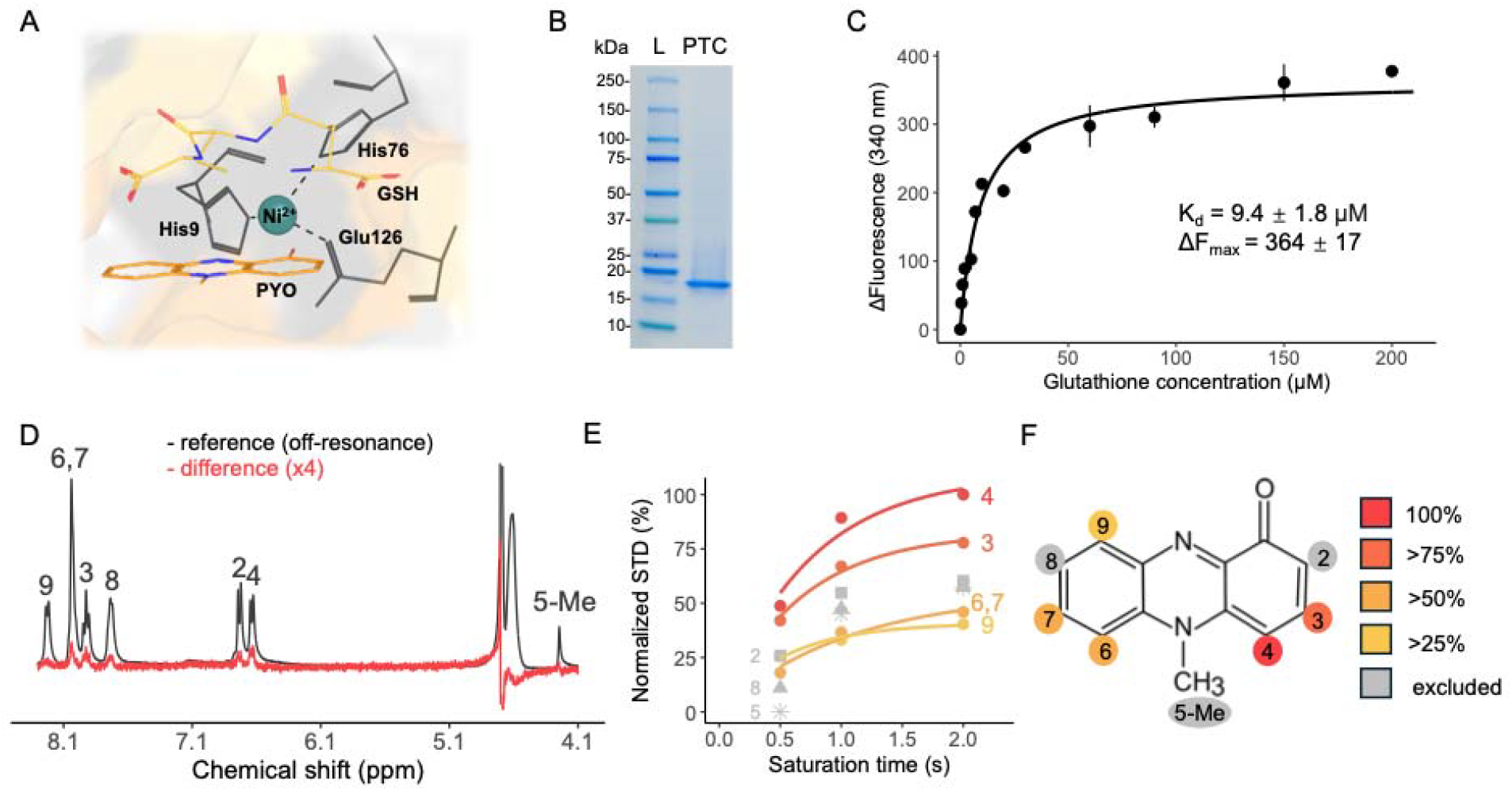
Identification, purification and characterization of the predicted phenazine-thiol conjugase. (A) Structural model of the protein, with an enlarged view of the active site showing glutathione and pyocyanin positioned in close spatial proximity to the divalent metal center. (B) SDS–PAGE analysis showing purified recombinant protein. L, ladder; PTC, the purified protein. (C) Hyperbolic fit used to determine the dissociation constant (K_d_) of the protein binding glutathione, determined by tryptophan fluorescence quenching. The error bar represents standard deviation of three replicates. (D) Reference ^1^H NMR spectrum of pyocyanin (black, off-resonance) and the STD NMR difference spectrum of the same sample, with (E) showing the STD build-up curve (STD intensity vs. saturation time) and (F) the mapped binding epitope of pyocyanin in its interaction with the protein.

Phenazine-thiol conjugase activity was assessed by incubating the purified enzyme with pyocyanin and freshly prepared reduced glutathione in the presence of various divalent metals. Catalytic activity was observed in reactions supplemented with Fe^2^□, Zn^2+^ or Ni^2^□, with the Ni^2^□-dependent reaction proceeding with the highest efficiency; subsequent experiments were therefore conducted under Ni^2^□-supplemented conditions. After 1.5 h of incubation, reaction mixtures exhibited a pronounced color change from blue (pyocyanin) to green (Figure 4A), whereas control reactions lacking enzyme remained blue. LC–MS analysis confirmed near-complete depletion of pyocyanin (*m*/*z* 211.0866 ± 10 ppm) and the formation of a new species at *m*/*z* 516.1548 ± 10 ppm, corresponding to a pyocyanin–glutathione conjugate ([C_23_H_25_N_5_O_7_S + H]^+^) (Figure 4B). We next tested 1-hydroxyphenazine as an additional phenazine substrate. Similar to pyocyanin, incubation with glutathione and the enzyme resulted in visible color changes (Figure 4C) and conversion of the parent compound (*m*/*z* 197.0710 ± 10 ppm) to a product with elemental composition [C□□H□□N□O□S + H]^+^ (*m*/*z* 516.1162 ± 10 ppm), consistent with a phenazine compound going through monooxygenation and glutathione conjugation (Figure 4D). Interestingly, for 1-hydroxyphenazine the enzyme accelerated formation of both conjugate isomers, whereas for pyocyanin it preferentially enhanced formation of a single isomer (Figure 4B,D). Moreover, the reaction with 1-hydroxyphenazine proceeded efficiently even in the absence of Ni^2^□, suggesting that our enzyme may have co-purified with other metals. Indeed, experimental characterization of the enzyme with ICP-MS revealed co-purification with Fe (38.4%), Ni (30.5%), Zn(25.9%) Mn (4.9%). These results indicate enzymatic flexibility in metal utilization depending on substrate chemistry, as observed for other VOC-family proteins (40). Intriguingly, neither phenazine-1-carboxylic acid (PCA) nor phenazine-1-carboxamide (PCN) showed detectable transformation under the same conditions as pyocyanin and 1-hydroxyphenazine, suggesting that an oxygen-containing group is a critical handle for enzymatic reactivity of PTC, consistent with the STD-NMR results indicating preferential binding to the oxygen-containing ring.

**Figure 4.**
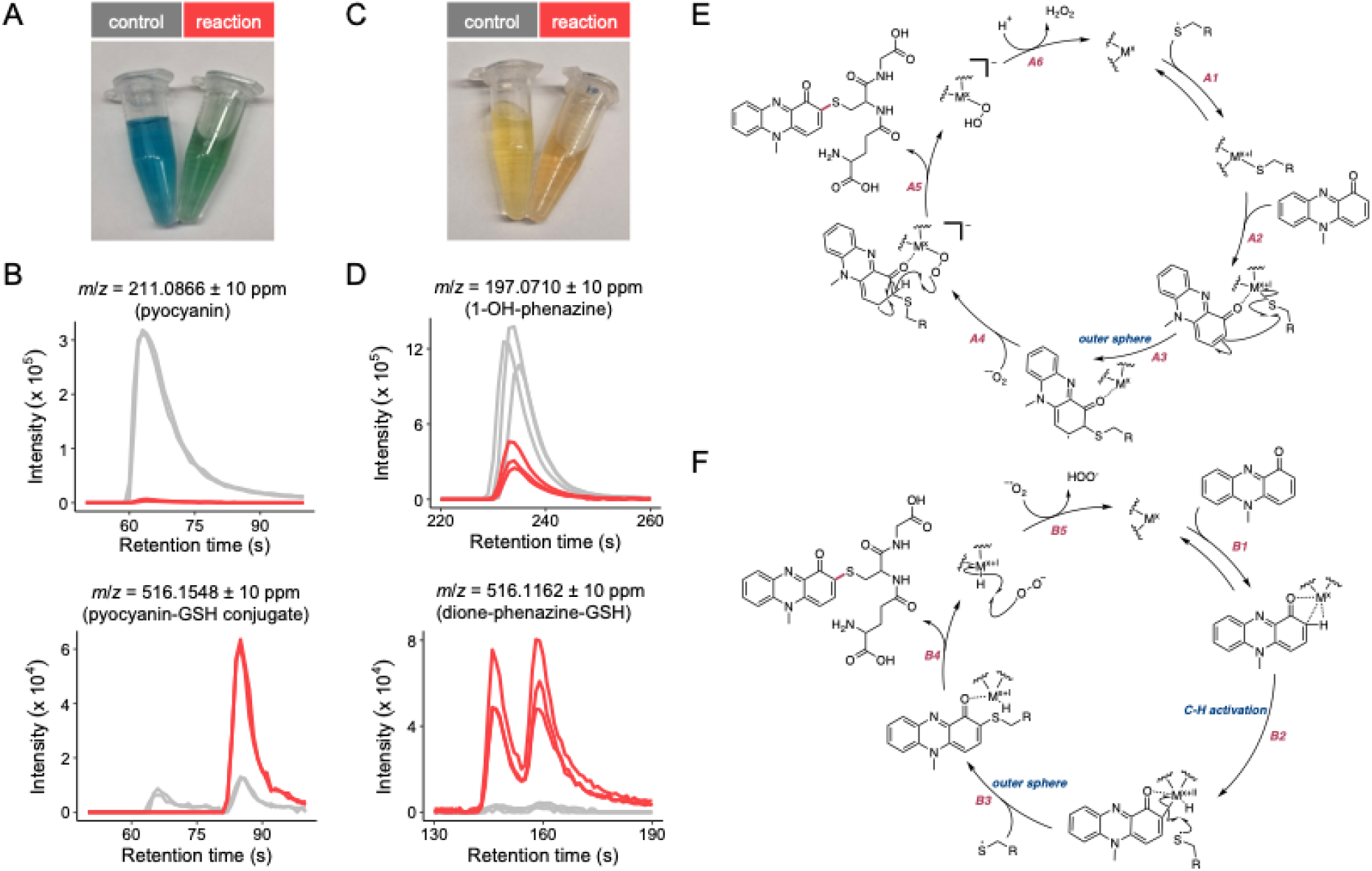
Determining the biochemical activities of phenazine-thiol conjugase (PTC). (A) Representative photographs of PTC reaction mixtures after 1.5 h incubation at room temperature with pyocyanin as substrate, compared to reactions with enzyme-free controls. (B) LC–MS chromatogram showing near-complete depletion of pyocyanin and formation of a new species with a m/z value consistent with a pyocyanin–glutathione conjugate. (C) Representative photographs of reaction mixtures using 1-hydroxyphenazine as substrate, with and without PTC. (D) LC–MS data showing consumption of 1-hydroxyphenazine and formation of a new species whose m/z is consistent with a product going through monooxygenation and glutathione conjugation. (E) Scheme A. The proposed conjugation mechanism that proceeds via oxidative addition. (F) Scheme B. The proposed conjugation mechanism that proceeds via C-H activation.

Two plausible enzyme mechanisms may explain the formation of the observed products. In one scenario (Scheme A, Figure 4E), the metal center may initially undergo oxidative addition via the GSH-based thiyl radical (Step A1), generated from the oxidation reaction between pyocyanin and GSH. This reaction would lead to the formation of a metal-sulfur bond and also superoxide (41, 42). Subsequently, the phenazine metabolite would coordinate to the metal (Step A2) and undergo outer sphere addition to the bound l-glutathione (Step A3) (43, 44). This species could stay ligated to the metal and superoxide could also coordinate, permitting a rapid hydrogen atom transfer (HAT) process, reestablishing aromaticity (Step A4) (45, 46). Following this, the presumed product would disengage the metal center (Step A5) and upon a proton transfer, hydrogen peroxide would be released as a byproduct (Step A6). Alternatively (Scheme B, Figure 4F), initial coordination of the carbonyl of the phenazine metabolite (Step B1), would be followed by C-H activation at the *ortho*-position of the ring (Step B2) (47, 48). This would then undergo an S_H_2 type reaction in which the thiyl radical could attack the phenazine metabolite in an outer sphere manner (Step B3), yielding the product and a metal hydride (Step B4), which could then be abstracted from the metal by superoxide to reestablish the active metal species (Step B5). Differentiating between these models will be the subject of future mechanistic studies.

More generally, our data indicate enzymatic catalysis of the following reactions:

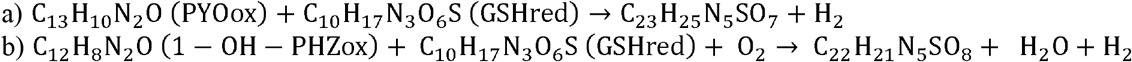

Accordingly, we designate the responsible enzyme <u>p</u>henazine-thiol <u>c</u>onjugase (PTC).

Beyond *P. agglomerans* where PTC was identified, we examined the distribution of similar enzymes in other bacteria. Screening the same set of > 200,000 non-redundant bacterial genomes (17), we identified 3,415 homologs of *ptc* (> 30% amino acid identity) spanning more than 30 phyla, including Pseudomonadota, Actinomycetota, and Acidobacteriota (Dataset S4). Genomes encoding *ptc* homologs were slightly enriched for phenazine producers compared with genomes lacking *ptc* homologs (0.51% vs. 0.15%; χ^2^ test, *P* = 1.15 × 10^-6^), suggesting a potential functional association between PTC-like enzymes and phenazine metabolism. Supporting this hypothesis, applying our model to this expanded homolog set, 2,423 sequences are predicted to potentially interact with phenazines (Dataset S4). To investigate the determinants underlying variation in predicted phenazine interactions, we assessed whether mutations near the phenazine-binding pocket exert a stronger influence on the model score than overall sequence divergence. We started by aligning all homologs (Methods) and identified residues within 5 Å of the bound phenazine for PTC. Because global sequence similarity itself affects the prediction score (Figure S3A), we regressed model scores for each PTC homolog against sequence identity to the *P. agglomerans* PTC and extracted the regression residuals, thereby isolating effects that transcend overall homology (Figure S3B). These detrended scores are no longer correlated with overall sequence homology, but exhibited a strong negative correlation with the number of amino acid substitutions within 5 Å of the bound phenazine (Pearson’s r = − 0.75, *P* = 0.002, Figure S3C), suggesting that mutations proximal to the ligand-binding site dominate predicted phenazine interactions relative to global sequence divergence.

Finally, we examined the physiological consequences of PTC-mediated phenazine thioconjugation using pyocyanin as a representative phenazine with well-documented antibiotic and virulence activities. Consistent with prior observations that thiol conjugation reduces phenazine toxicity (35), reaction products generated by PTC displayed substantially reduced toxicity toward *P. agglomerans* compared to unreacted pyocyanin (Figure 5A). Similar results were observed for the distantly related, phenazine-sensitive bacterium *Chryseobacterium shigense*, indicating broad relevance of the detoxifying effect (Figure 5A). Given that the thiolated product of PTC is less toxic than pyocyanin, we hypothesized that deletion of the gene encoding PTC in *P. agglomerans* would result in a fitness defect under pyocyanin stress. However, in comparing the growth of wild-type and Δ*ptc* strains in the presence of 100 µM pyocyanin, no significant growth difference was observed (Figure 5B). At higher pyocyanin concentrations (400 µM), however, the wild-type strain exhibited a growth disadvantage relative to the Δ*ptc* strain (Figure 5B), potentially due to excessive consumption of intracellular thiol compounds. Consistent with this notion, supplementation of the growth medium with glutathione restored similar growth levels in the wild-type and Δ*ptc* strains, and the observed phenotypes were further validated by genetic complementation of *ptc* (Figure S4). Together, these results raise intriguing questions regarding the physiological role of PTC within *P. agglomerans*, motivating future studies. These results also underscore that the unexpected phenotypic effects would likely have precluded discovery of this enzyme through screens scoring for loss of phenazine resistance, highlighting the value of phenotype-independent, sequence–driven approaches for targeted discovery of antibiotic-modifying enzymes that may be leveraged for biocatalysis.

**Figure 5.**
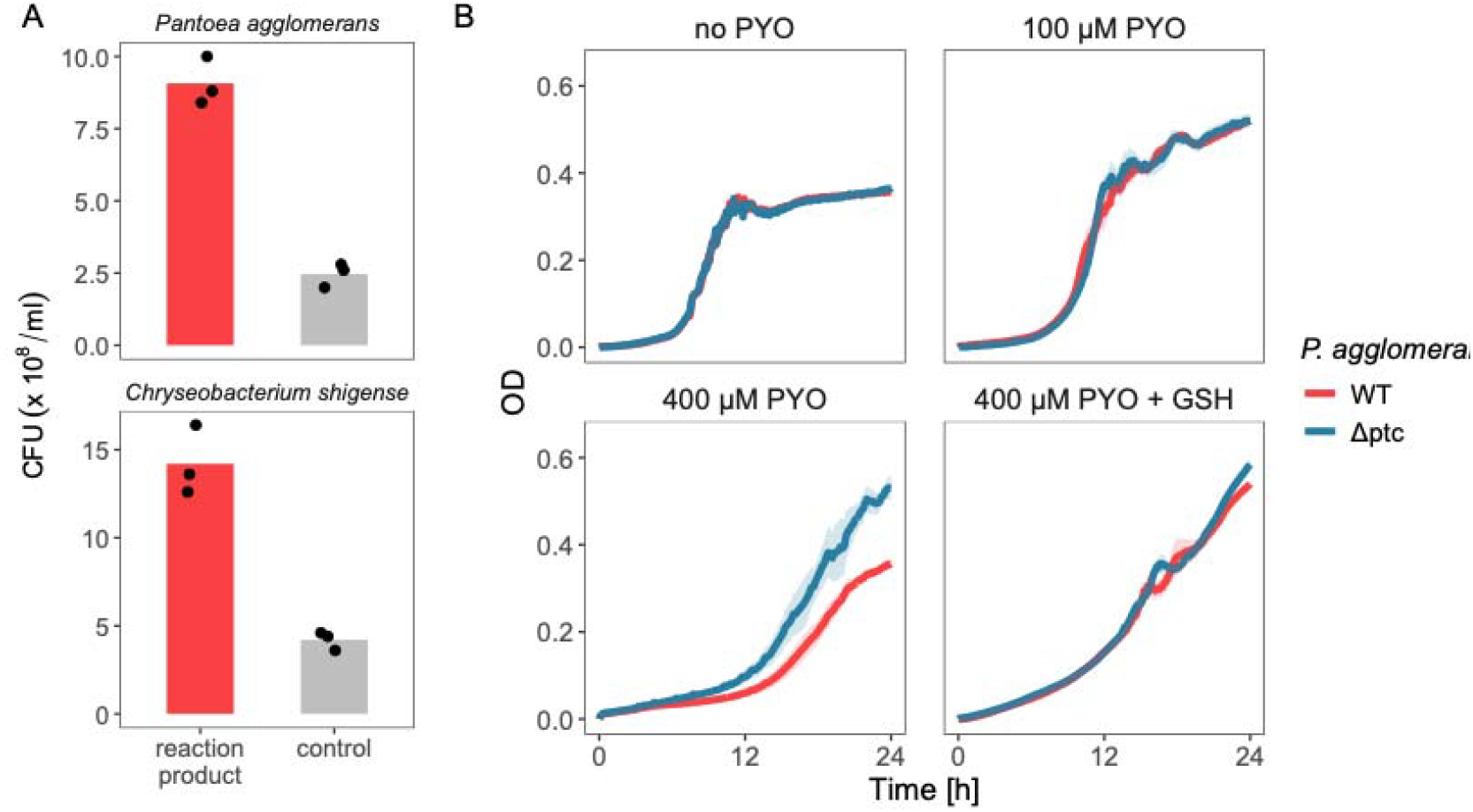
Physiological consequences of PTC-mediated phenazine thioconjugation. (A) Thioconjugated pyocyanin generated by PTC exhibits reduced toxicity compared with unreacted pyocyanin as a control in both *Pantoea agglomerans* and the distantly related phenazine-sensitive bacterium *Chryseobacterium shigense*. (B) The Δ*ptc* mutant shows no growth disadvantage compared to the wild-type in the presence of 100 µM pyocyanin. It displays a growth advantage over the wild-type strain at 400 µM pyocyanin, possibly because enzymatic thioconjugation depletes intracellular thiol pools, imposing additional stress. Supplementation with glutathione restores comparable growth between wild-type and Δ*ptc* strains.

## Discussion

Using the modification of phenazine antibiotics as a case study, we have demonstrated that it is possible to merge comparative genomics with protein machine learning to develop a simple yet generalizable framework for zero-shot identification of enzymes acting on data-sparse substrates. Starting from only 14 known phenazine-modifying enzymes, we show that genome-informed data augmentation can convert extremely limited biological knowledge typically unsuitable for machine learning into actionable predictive power. The general pipeline we have developed has the potential to accelerate solutions to many biological problems that lack the scale of curated data typically required for machine learning. The same logic can be generalized to other systems, as long as (1) conserved genomic context suggests informative functional labels (e.g., polysaccharide utilization loci, biosynthetic gene clusters for metabolites, phage defense islands, and conserved operons underlying various metabolic functions) and (2) the target activity leaves a learnable imprint on protein sequence (e.g., substrate binding, cellular localization, optimal biophysical conditions).

Using this genome-informed learning framework, we identified PTC, to our knowledge the first enzyme shown to catalyze phenazine–thiol conjugation. Although thiol-conjugated phenazines have been detected for decades, these reactions were presumed to arise from spontaneous chemical reactivity. The identification of PTC establishes that phenazine thioconjugation can be enzymatically catalyzed, revealing a regulated biochemical mechanism for modulating phenazine toxicity rather than an uncontrolled detoxification byproduct. Canonically, glutathione conjugation to toxic or electrophilic compounds is often mediated by glutathione S-transferases (49), making the discovery of a glyoxalase-type enzyme catalyzing phenazine–thiol conjugation both unexpected and mechanistically intriguing. This finding raises new questions regarding the catalytic flexibility of glyoxalase-like enzymes, including their substrate scope and flexible metal dependence. Our structural modeling, ligand-binding analyses, and product characterization together provide a foundation for future mechanistic dissection of this reaction.

That phenazine-thiol conjugation yields a product that is less toxic than its substrate, but shows no demonstrable fitness advantage to the organism catalyzing this reaction (and possibly even a disadvantage, if intracellular glutathione pools are severely depleted) is intriguing. Because growth arrest is known to promote tolerance to antibiotics (50), it is conceivable that PTC may facilitate tolerance to phenazines. On a larger scale, it is possible that organisms with PTC may stabilize the microbial communities of which they are members. These, and other hypotheses, await future investigations. Beyond bacterial physiology, we note that glutathione is a central metabolite across domains of life (51, 52), with dysregulated glutathione homeostasis implicated in diverse human diseases, including cancers where elevated glutathione levels contribute to chemoresistance (53). Phenazines are known to perturb intracellular glutathione pools (54). Accordingly, elucidating the enzymatic mechanisms governing phenazine–glutathione interactions has implications that extend beyond bacterial secondary metabolism, with the potential to inform our understanding of host–pathogen interactions and redox balance in human disease.

## Methods

### Data collection

To assemble a high-quality training set of phenazine-modifying enzymes, we curated 14 unique genes for which experimental evidence of modifying phenazines is available in the literature. We restricted our dataset to enzymes that act directly on the phenazines as the substrate to maintain a conservative positive training set. These 14 genes exhibit no detectable homology to each other, while each of them is represented by one to several homologs from different bacterial taxa, mostly in the vicinity of the phenazine biosynthesis genes. Details of these 14 genes are provided in Table S1.

To identify phenazine producers, we searched for genomes containing homologs of the canonical *phzABCDEFG* gene cluster as previously described (16). Given that *phzA* and *phzB* are homologous to each other, and *phzC* is often functionally promiscuous, we required that genomes contain at least five of the seven genes, located within a continuous window of ten genes. Using *Pseudomonas aeruginosa* PA14 *phzA–G* sequences as references, we computationally screened 202,601 bacterial genomes from the GlobDB collection (v220; accessed October 2, 2024) (17). These genomes are non-redundant, each representing a unique species-level taxon. Our search yielded 310 different phenazine producers across the Bacteria domain. An additional set of 7 phenazine producer genomes identified via NCBI BLAST but not present in GlobDB were manually added, resulting in 317 unique phenazine producers. Across these producers, homologs of the 14 modifying genes were identified using MMseqs2 (55) with a cutoff of 50% coverage and 25% amino acid identity. Among them, those that are located within ten genes of the *phzA– G* core cluster were labelled as positive. We adopted here a cut-off of 10 genes because some characterized phenazine biosynthesis gene clusters (e.g., *ehpA-O*) spans as far as 10 genes beyond the core *phzA-G* genes (56). Despite this permissive cutoff, 91.8% of identified positives were located fewer than 5 genes from the core cluster, reflecting strongly conserved genomic organization. After dereplication at 90% sequence identity, this procedure yielded 593 positive sequences from 239 different phenazine producers. Homologs located far from the core cluster (>50 genes away) were designated as “hard negatives” for evaluation (Figure 2C–E) to deliberately challenge the model, although we note that true phenazine-modifying enzymes may also reside outside biosynthetic clusters (e.g., *phzH* in *P. aeruginosa*).

We built a balanced set of negative sequences to enable the contrastive learning framework. We collected seed negative sequences from reviewed, unfragmented UniProt entries (accessed November 20, 2025) belonging to the same functional categories as the positive sequences (i.e., amidotransferase, methyltransferase, monooxygenase, prenyltransferase, AMP-binding enzymes and dehydrogenase), but acting on well-characterized substrates irrelevant to phenazines such as DNA, RNA and amino acids. A full list of seed negative sequences is provided in Table S2. Because reviewed UniProt entries exhibit strong taxonomic bias toward model microorganisms, directly mixing these sequences with positives risks training a classifier that merely captures phylogenetic signatures. To mitigate this, UniProt sequences were used only as seed references, and their homologs were then searched within the same 239 phenazine-producer genomes that contain modifying genes used as positive sequences. This ensures shared evolutionary backgrounds between positive and negative sequences. After dereplication at 80% sequence identity, this procedure yielded 589 negative sequences balancing the number of positive sequences in each of the six functional categories (Table S3).

We identified PTC homologs also from the same GlobDB genome collection using MMseqs2 (>30% sequence identity, >80% coverage) (55). To efficiently construct a high-quality multiple sequence alignment for >3000 sequences, we adopted a centroid-based clustering strategy as developed before (57). Specifically, sequences were clustered with MMseqs2 (-c 0.5 --min-seq-id 0.5) such that each cluster represented sequences with ≥50% pairwise identity. This reduced the dataset to 135 representative (centroid) sequences. These centroid sequences were aligned with MAFFT (version 7.505, --maxiterate 10 --localpair) (58) to generate a backbone alignment. The remaining sequences were subsequently added to this alignment using the --addfragments option in MAFFT.

### Model architecture

All protein sequences were embedded using the ESM Cambrian 600M pretrained protein language model (23), yielding 1,152-dimensional embeddings per sequence. Mean-pooled embeddings were passed into a three-layer multilayer perceptron (MLP) with 512, 128 and 64 dimensions with ReLU activations and dropout of 0.1. The MLP was trained using contrastive learning to bring phenazine-interacting enzymes closer in the latent space while separating them from non-interacting enzymes. Given the learnt embeddings from protein language models have learnt patterns across the protein landscape, the largest signals typically correspond to broad functional classes. As our objective was to learn a specific transformation (phenazine-modification) which spans multiple broad classes, we designed a pair scheme for this purpose. To that end, we constructed non-redundant sequence pairs belonging to four categories: (1) a positive and a negative sequence from the same functional category (e.g., a phenazine-modifying versus adenine-modifying methyltransferase), (2) two positive sequences from different functional categories (e.g., a phenazine-modifying methyltransferase and monooxygenase), (3) two negative sequences from different functional categories, and (4) a positive and a negative sequence from different functional categories. Pairs from categories (2) and (3) were labeled as positive pairs, whereas pairs from categories (1) and (4) were labeled as negative pairs. The numbers of pairs sampled from categories (1)– (4) were 71,904 (exhaustive), 60,000, 1,000, and 1,000, respectively, totaling 133,904 with heavier sampling of categories (1) and (2) to emphasize learning features that distinguish phenazine-interacting enzymes from non-interacting homologs within the same functional classes. Contrastive learning was performed using a margin-based contrastive loss defined on the Euclidean distance d between latent embeddings: the loss was set to d^2^ for positive pairs and max{0,(m-d)^2^} for negative pairs, with a margin m = 1. A scaling factor of 1,000 was applied to stabilize the loss. (59)The contrastively learned latent embeddings were subsequently used to train a support vector machine classifier (linear kernel, C = 1.0, class-weight balanced) to distinguish phenazine-modifying enzymes from non-modifying ones. The models were implemented in Python using PyTorch (60). Model performance was evaluated using balanced accuracy and area under the precision–recall curve (AUPR), with balanced accuracy defined as the mean of sensitivity and specificity. For evaluation on hard-negative test sets (Figure 2C–E), AUPR was reported relative to a baseline AUPR, defined as the fraction of positive sequences in the test set to account for the null expectation under class imbalance. Model prediction scores for all proteins encoded in the *Pantoea agglomerans* genome are provided in Dataset S3. Training was performed using AdamW (49) with a learning rate of 0.0001, batch size of 512, and 10 epochs.

### Protein expression and purification

The PTC expression construct was generated using Sequence and Ligation Independent Cloning (SLIC) as described by (61). Primers 3C-LP1 and ccdB-LP2 (Table S5) were designed to introduce a human rhinovirus (HRV) 3C protease cleavage site and a ccdB counterselection cassette (61). The PTC coding sequence was amplified by PCR using KAPA HiFi DNA polymerase from genomic DNA isolated from *Pantoea agglomerans* W2I1 (NCBI assembly GCF_030815835.1) (DNeasy Blood & Tissue Kit, Qiagen). The PCR product was assembled into the pCoofy18 vector, which encodes an N-terminal His□□ tag, using the Gibson Assembly® Cloning Kit (New England Biolabs).

The resulting plasmid (N-His□□-PTC) was transformed into *Escherichia coli* BL21(DE3) for protein production. Cultures were grown in LB medium supplemented with 50 mg/L kanamycin at 37 °C with shaking at 180 rpm. Induction with IPTG was omitted, as higher protein yields were obtained from uninduced cultures, likely due to leaky expression from the T7 promoter. Cells were harvested 6 h after cultures reached an OD□□□ of 0.6 and stored at –80 °C.

For protein purification, frozen cell pellets were thawed and resuspended in lysis buffer 50 mM Tris-HCl, pH 7.6, 250 mM NaCl supplemented with protease inhibitor cocktail (Roche) and DNase I (Roche). Cells were lysed using an Avestin Emulsiflex C3 homogenizer (three passes at 15,000 psi). The lysate was clarified by ultracentrifugation at 42,000 rpm for 50 min at 4 °C. The supernatant was applied to a HisTrap™ affinity column (Cytiva), and PTC was purified by stepwise elution with increasing concentrations of imidazole in the same buffer. PTC eluted at imidazole concentrations above 300 mM. Eluted fractions were analyzed by SDS–PAGE, followed by BlueSafe staining (NZYTech) and His-tag– specific staining (InVision™ His-tag) to confirm purity and identity. Fractions containing pure PTP were pooled, concentrated, and buffer-exchanged using Amicon® Ultra centrifugal filters (Millipore; 10-kDa MWCO). Protein concentration was determined by absorbance at 280 nm using a theoretical extinction coefficient of 16,690 M□^1^ cm□^1^ calculated with ExPASy ProtParam.

### Modeling protein structure and ligand binding

PTC was initially modelled using AlphaFill (36) with the UniProt reference sequence A0A379LSW4 as input. The resulting model showed approximately 30% sequence identity to experimentally solved glyoxalase I structures from multiple organisms deposited in the Protein Data Bank (PDB IDs: 1QIP, 4MTQ, 1FRO, and 1BH5), including human, murine, and bacterial homologs. Ligands were extracted from templates exhibiting high template confidence scores (TCS > 0.8) and low local root-mean-square deviation values (l-RMSD < 0.5 Å), including glutathione from 1QIP chain B, Ni^2^□ from 4MTQ chain B, and Zn^2^□ from 1FRO chain A and 1BH5 chain D. All template structures corresponded to glyoxalase I proteins, which adopt a conserved homodimeric architecture; therefore, all subsequent modeling was performed assuming a dimeric quaternary structure.

Prediction of ligand-bound conformations was performed using Chai-1 (37) with the full-length protein sequence extended by the polyhistidine tag and HRV 3C protease cleavage site. Ligands were provided as SMILES strings obtained from PubChem: pyocyanin (CN1C2=CC=CC=C2N=C3C1=CC=CC3=O), 1-hydroxyphenazine (C1=CC=C2C(=C1)N=C3C=CC=C(C3=N2)O), Glutathione reduced (C(CC(=O)N[C@@H](CS)C(=O)NCC(=O)O)[C@@H](C(=O)O)N), Ni^2^□ ([Ni+2]), Fe^2^□ ([Fe+2]), Mn^2^□ ([Mn+2]), and molecular oxygen (O=O). Molecular visualizations were generated using the PyMOL Molecular Graphics System (Version 3.0 Schrödinger, LLC.).

### Standard enzymatic assays

Standard reactions contained 10 µM PTC, 100 µM of the indicated phenazine, and 10 mM reduced glutathione (freshly prepared immediately before use). For assays examining pyocyanin–glutathione conjugation, NiCl□ was additionally included at a final concentration of 500 µM.

For LC-MS analysis, reactions were incubated at room temperature in microcentrifuge tubes protected from light (wrapped in aluminum foil) for approximately 1.5 h before filtering the protein to stop the reaction using a VWR centrifugal filter with 10KDa cutoff.

For kinetic measurements, reactions were instead aliquoted into three wells of a 96-well plate (GenClone**®**), and reaction progress was monitored by measuring absorbance at 450 nm, corresponding to the characteristic signal of the pyocyanin–glutathione conjugate.

### LS-MS measurements

For LC–MS analysis, all samples were diluted 10-fold with water. Analyses were performed on an Agilent 1200 series system configured with a G1312B binary pump connected to an Agilent 6230 time of flight (TOF) mass spectrometer equipped with a JetStream electrospray ionization source. Chromatographic separations were performed on a ZORBAX Extend-C18 column (Agilent) with dimensions 2.1 x 50 mm and a particle size of 1.8 µm. The column temperature was 30 °C and the flow rate was 0.5 mL/min. Mobile phase A was water with 0.025% acetic acid and mobile phase B was acetonitrile. The separation used a linear gradient from 2% to 95% B over 0–10 min, 95% B over 10–11 min, and a re-equilibration at 2% B over 11–13 min. The injection volume was 1 µl, and the first 30 seconds were diverted to waste to avoid overloading the MS detector. For detection, the MS parameters were: gas temp, 320 °C; drying gas, 8 L/min; nebulizer, 35 psi; sheath gas temp, 325 °C; sheath gas flow, 11 L/min; fragmentor, 175 V; skimmer, 65 V; Oct 1 RF Vpp, 750 V; scan time, 1s. A calibration solution was infused with reference masses set to 121.0509 and 922.0098 in the positive ionization mode.

### STD-NMR experiments

Saturation transfer difference (STD) NMR experiments were performed on a Bruker Avance III 400 MHz spectrometer equipped with a broadband iProbe, following a protocol adapted from (62). Samples contained 2 mM pyocyanin and 60 μM PTC in 50 mM phosphate buffer (pH 7.6), prepared in D□O. All experiments were conducted at 298 K.

STD spectra were acquired using the Bruker pulse program stddiffgp19, with 256 transients, 16k data points in the direct dimension, and a spectral width of 15.6191 ppm centered at 4.703 ppm to ensure efficient water suppression. Selective saturation was applied within the aliphatic protein resonance region, with the on-resonance frequency set to 0 ppm and the off-resonance frequency set to 40 ppm, using a saturation power of 10 μW. Control experiments performed in the absence of protein confirmed the absence of direct ligand saturation under these conditions. STD buildup curves were recorded at saturation times of 0.5, 1.0, and 2.0 s. STD intensities were calculated according to

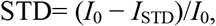

where *I*_0_ and *I*_STD_ correspond to peak intensities in the off-resonance and STD spectra, respectively.STD buildup curves were analyzed quantitatively by fitting the STD intensity as a function of saturation time to a monoexponential model,

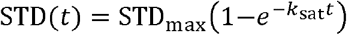

using nonlinear least-squares regression implemented in Python (SciPy). For each ligand proton, the fitted parameters STD_max_ and *k*_sat_were extracted together with their standard errors derived from the covariance matrix of the fit. To ensure robustness and reproducibility, only fits meeting predefined quality criteria were retained for further analysis: relative uncertainty ≤30% for STD_max_ and ≤50% for *k*_sat_. Protons failing to meet these criteria were excluded from epitope mapping. Following filtering, STD_max_ values were normalized post-fit to the largest retained STD_max_ and expressed as percentages. The initial slope (STD_max_ · *k*_sat_) was additionally calculated as a measure of early saturation transfer efficiency.

### Tryptophan quenching

All quenching experiments were performed using a Molecular Devices SpectraMax M3 Multi-Mode spectrophotometer using 3 mL quartz cuvette with 1cm pathlength.

For tryptophan quenching the excitation wavelength was fixed to 290nm and emission spectra were collected between 310 and 400nm with a slit width of 9nm. Temperature was maintained at 25□°C, and samples were gently resuspended three times after each ligand addition before measurement. To measure PTC–glutathione (reduced) interactions, 1□µM PTC was titrated with increasing concentrations of glutathione (0–200□µM) freshly prepared in 50□mM Tris, pH□7.6, 250□mM NaCl, the same buffer used for all titrations. Fluorescence intensity at 340□nm from two replicate measurements was corrected for inner-filter effects. The resulting change in fluorescence (ΔF) was fitted directly using a hyperbolic binding model:

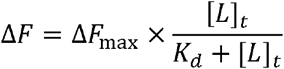

where [*L*]_t_ is the total ligand concentration, *K*_*d*_is the apparent dissociation constant, and Δ*F*_max_is the fluorescence change at saturation. This model is appropriate for these conditions because the protein concentration (1□µM) is much lower than the observed *K*_*d*_ (∼10□µM), so ligand depletion is negligible.

### ICP-MS measurements

Concentrations of cobalt, chromium, copper, iron, manganese, nickel, vanadium and zinc were determined by Inductively Coupled Plasma – Mass Spectrometry (ICP-MS) using an Agilent 8800. 50 μL of each protein sample (duplicates) was diluted to 15 mL with 5% nitric acid. The sample introduction system consisted of a Miramist nebulizer, Scott type spray chamber and fixed injector quartz torch. A guard electrode was used, and the plasma was operated at 1500 W. All elements were determined in helium MS-MS mode. Standards were prepared from an ICP-MS standard mix of first row transition elements (CCS – 6, Inorganic Ventures, Christiansburg, VA, USA) in the range of 0 to 400 μg/L of each element. A calibration verification (CV) solution was prepared from multielement mix p/n 044518 from Thermo Scientific. Agreement between standards and the CV was better than 4% for all metals. Drift during the measurement time was less than 4% for all metals.

### Gene deletion and complementation

Gene deletion of the *ptc* gene in *Pantoea agglomerans* was performed using a modified λ-Red recombineering protocol (63, 64). Approximately 500 bp genomic regions upstream and downstream of *ptc* were PCR-amplified and purified by gel extraction (Monarch #T1120L). The kanamycin resistance cassette from plasmid pKD4 was similarly amplified, with 50-bp homologous overhangs added to each end to overlap with the upstream and downstream fragments. The three fragments (upstream arm, kanamycin cassette, downstream arm) were fused into a single linear deletion construct by overlap-extension PCR, first 15 rounds using annealing temperature of the overhang region then 30 rounds using the annealing temperature of the primers. Plasmid pKD78, which carries the λ-Red recombination system and chloramphenicol resistance marker, was first introduced into the strain by electroporation (1.8 kV, 25 μF, 200 Ω, 1-mm cuvette), followed by recovery in LB at 30 °C for 1 h and selection on LB agar supplemented with 12.5 μg/mL chloramphenicol. Verified pKD78-containing transformants were grown in LB with 0.2% arabinose and chloramphenicol until OD ≈ 0.6, harvested, and washed twice with ice-cold glycerol (2,000 × g, 10 min) to prepare electrocompetent cells. The fused linear deletion fragment was electroporated into these induced competent cells using the same electroporation settings, and cells were recovered and selected on LB agar containing 50 μg/mL kanamycin. Candidate deletion mutants were screened by colony PCR, and confirmed colonies were streaked on LB agar at 39 °C to cure pKD78; colonies losing chloramphenicol resistance were subsequently transformed with plasmid pCP20 with chloramphenicol resistance marker to excise the kanamycin cassette via FLP recombination. After a second curing step at 39 °C to remove pCP20 and screening for the loss of chloramphenicol resistance, the final unmarked *ptc* clean deletion mutant was verified by PCR genotyping and genome sequencing.

The Δptc mutant was complemented using the broad-host-range plasmid pBBR1MCS-2. The ptc coding sequence was amplified by PCR from genomic DNA isolated from *Pantoea agglomerans* strain W2I1 (DSM 3493) using KAPA HiFi DNA polymerase (Roche) and the DNeasy Blood & Tissue Kit (Qiagen). PCR primers are listed in Table S5. The resulting amplicon was cloned into the pBBR1MCS-2 vector, which carries a kanamycin resistance marker, using Gibson Assembly (New England Biolabs) according to the manufacturer’s instructions. The assembled plasmid was introduced into *P. agglomerans* by electroporation as described above.

### Bacterial survival assays with modified phenazines

Wild-type strains of *Pantoea agglomerans* W2I1 and *Chryseobacterium shigense* W4I1 were grown overnight with shaking in LB medium at 30°C. Phenazine reactions were performed as described above, except that the indicated phenazine was used at a final concentration of 400 µM and reaction mixtures were sterile-filtered to prevent contamination. Reactions were mixed 1:1 (v/v) with bacterial cultures adjusted to an OD□□□ of 1.0 (100 µL reaction + 100 µL cells in LB), resulting in a final OD□□□ of 0.5 in diluted LB medium. Mixtures were incubated with shaking overnight at 30°C in 96-well plates. Following incubation, cultures were serially diluted and plated onto LB agar. Colonies were grown for 24 h at 30°C and quantified to assess survival.

### Culture conditions

Strains were grown overnight with shaking in LB medium at 30 °C, then diluted to a starting OD□□□ of 0.025 in Rhizosphere Medium (RZM). RZM was formulated to approximate the ionic composition and carbon availability of wheat rhizosphere environments. The base medium contained MgCl□·6H□O (0.406 mM), CaCl□·2H□O (0.68 mM), KCl (6.71 mM), phosphate (2 mM total, pH 7.0), ammonium nitrate (15 mM) as a nitrogen source, and was buffered with MES (50 mM, pH 6.0). D-glucose (20 mM) was provided as the sole carbon source. Trace elements were included at final concentrations of FeSO□·H□O (50 µM), MnSO□·H□O (50 µM), CoCl□·6H□O (52.5 nM), CuSO□·5H□O (50 nM), ZnSO□·7H□O (15 µM), Na□MoO□·2H□O (0.5 µM), potassium iodide (2.5 µM), Na□SeO□ (10 nM), NiSO□·6H□O (1.0 µM), H□BO□ (50 µM), and EDTA (275 µM). Vitamins were supplied by addition of the ATCC Vitamin Supplement (MD-VS), which provides folic acid, biotin, pyridoxine, thiamine, riboflavin, nicotinic acid, calcium pantothenate, vitamin B□□, p-aminobenzoic acid, and thioctic acid at concentrations based on Wolfe’s vitamin solution. The complete medium was prepared in Milli-Q water and sterilized by filtration. Phenazines were added where indicated; pyocyanin was synthesized as described previously (65) and 1-hydroxyphenazine was purchased from Sigma-Aldrich.

## Supporting information

SI datafiles

## Code availability

The codes of ML-CITO are publicly available at https://github.com/Xiaoyu2425/MLCITO/blob/main/ML-CITO_Pantoea_agglomerans.ipynb.

## Data availability

All data used in this study has been provided in the paper or supplementary information.

## Acknowledgement

We thank Ricardo O. Louro, Yueming Long, Frances Arnold, Otto Cordero, Lev Tsypin, Jinyang Li, Reinaldo Alcalde, Calvin Rusley and all members of the Newman lab for helpful discussions. We thank Scott C. Virgil for support with the mass spectrometry and Nathan Dalleska for the ICP-MS analysis. The ICP-MS equipment is located in the Water and Environment Laboratory at the California Institute of Technology and is supported by the Resnick Sustainability Institute. X.S thanks Yunha Hwang for illuminating discussions on protein language models. X.S thanks Ming Chen, Jeffery Barrick and Devin Lake for generously sharing protocols and materials for genetics. Grants from the NIH (2R01AI127850-06A1) to DKN, EMBO (ALTF 191-2023) to IBT, and a Nemko Postdoctoral Fellowship (to XS) from the Division of Biology and Biological Engineering supported our research. AM was supported by the Schmidt Science Fellows, in partnership with the Rhodes Trust. SJC is grateful to Michael and Alice Jung for endowing the Jung Chair in Medicinal Chemistry and Drug Discovery at UCLA, which partially supported this work.

## Notes

### Competing Interest Statement

The authors have declared no competing interest.

