## Supporting Information for "Discovery of a phenazine–thiol conjugase from sparse data using genome-informed machine learning"

**This PDF file includes:**

Figures S1 to S4

Tables S1 to S5

Legends for Datasets S1 to S4

**Other supporting materials for this manuscript include the following:**

Datasets S1 to S4

**Figure S1** Schematic illustration of “hard negative” test in Figure 2C-D.

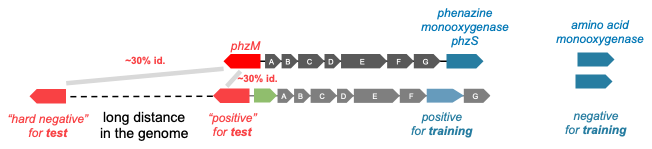

**Figure S2** Tryptophan fluorescence quenching spectra of PTC upon addition of glutathione.

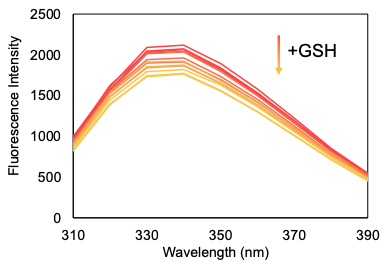

**Figure S3** Binding-site variation affects model scoring beyond global sequence homology.

(A) Relationship between model score and amino acid sequence identity to Pantoea agglomerans PTC across 3,415 PTC homologs, showing a positive correlation driven by overall sequence similarity. (B) Model scores after detrending for global sequence homology by extracting the regression residual of the model score, demonstrating removal of homology-dependent effects. (C) Detrended model scores plotted against the number of amino acid substitutions within 5 Å of the bound phenazine, revealing a significant negative correlation, indicating that mutations proximal to the ligand-binding site disproportionately reduce predicted phenazine interaction independent of overall sequence divergence. Error bars denote ± SEM.

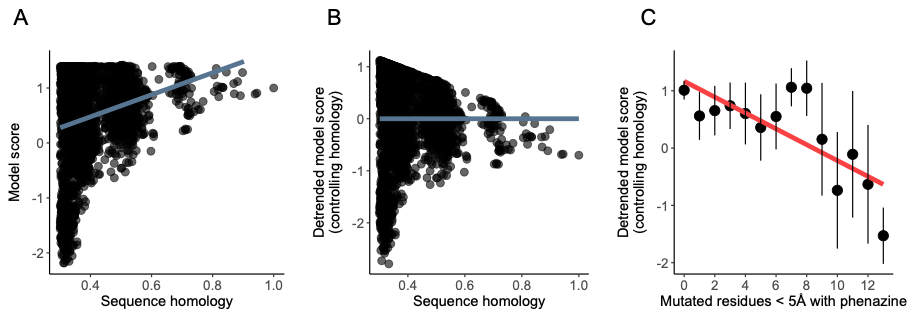

**Figure S4** Growth phenotype of the wild-type (WT), *ptc* deletion strain (Δ*ptc*), and the genetic complementation strain (Δ*ptc*+pBBR1MCS-2::*ptc*). The *ptc* complementation strain exhibits growth deficit at high concentration of pyocyanin, which can be rescued by adding excessive amount of glutathione.

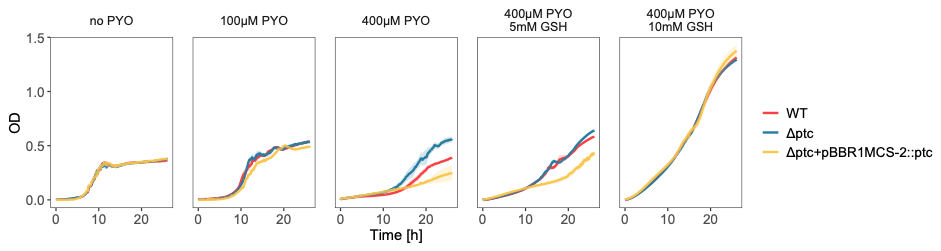

Table S1 Detailed information of the seed positive enzymes known to modify phenazines.

| **homologous group ID** | **representative** | **functional category** | **UniProt entry** | **other reported homologs** |
| --- | --- | --- | --- | --- |
| G01 | phzH | amidotransferase | Q9I781 | / |
| G02 | phzM | methyltransferase | Q9HWH2 | LaPhzM, ppzM, epzM |
| G03 | pcm2 | methyltransferase | A0AAQ0BTI5 | lomo11 |
| G04 | ehpI | methyltransferase | O32599 | xpzL |
| G05 | phzS | monooxygenase | Q9HWG9 | dapS, xpzG |
| G06 | phzO | monooxygenase | E3UVD3 | / |
| G07 | LaPhzNO1 | monooxygenase | A0A172J1R7 | NaPhzNO1, xpzH |
| G08 | lomo10 | monooxygenase | A0A0E3UR82 | / |
| G09 | mpz10 | prenyltransferase | A0A068CM06 | CnqPT1 |
| G10 | ppzP | prenyltransferase | C4PWA1 | epzP |
| G11 | ehpF | AMP-binding adenylation | Q8GPH0 | esmB1, lomo15 |
| G12 | ehpG | dehydrogenase | Q8GPG9 | esmB2, lomo16 |
| G13 | ehpL | dehydrogenase | Q8GPG8 | pcm1, lomo9 |
| G14 | ehpK | dehydrogenase | O32601 | xpzN |

Table S2 Detailed information of the seed negative enzymes from the same broad functional categories with positive sequences.

| homologous group seed ID | functional category | Uniprot entry |
| --- | --- | --- |
| NG01 | AMP-binding enzyme | Q9Z3R3 |
| NG02 | AMP-binding enzyme | P27550 |
| NG03 | AMP-binding enzyme | Q88DW6 |
| NG04 | AMP-binding enzyme | P19409 |
| NG05 | AMP-binding enzyme | A1A787 |
| NG06 | AMP-binding enzyme | B2HKM1 |
| NG07 | AMP-binding enzyme | P0DX14 |
| NG08 | AMP-binding enzyme | P18204 |
| NG09 | AMP-binding enzyme | A0R618 |
| NG10 | AMP-binding enzyme | B2HIN2 |
| NG11 | AMP-binding enzyme | P9WQ47 |
| NG12 | AMP-binding enzyme | Q02278 |
| NG13 | AMP-binding enzyme | O33855 |
| NG14 | AMP-binding enzyme | P38135 |
| NG15 | AMP-binding enzyme | P55912 |
| NG16 | amidotransferase | A0QUW9 |
| NG17 | amidotransferase | A1R4R2 |
| NG18 | amidotransferase | P04079 |
| NG19 | amidotransferase | P17169 |
| NG20 | amidotransferase | A0L5G0 |
| NG21 | amidotransferase | P58251 |
| NG22 | amidotransferase | Q8Y489 |
| NG23 | amidotransferase | P39648 |
| NG24 | dehydrogenase | P37769 |
| NG25 | dehydrogenase | E5KIB9 |
| NG26 | dehydrogenase | P52643 |
| NG27 | dehydrogenase | P0AEK7 |
| NG28 | dehydrogenase | P11886 |
| NG29 | dehydrogenase | O34214 |
| NG30 | dehydrogenase | P0AC53 |
| NG31 | dehydrogenase | P54226 |
| NG32 | dehydrogenase | P9WN81 |
| NG33 | dehydrogenase | Q3K4Z1 |
| NG34 | dehydrogenase | P61889 |
| NG35 | dehydrogenase | Q9RCG0 |
| NG36 | dehydrogenase | P00370 |
| NG37 | dehydrogenase | P07003 |
| NG38 | dehydrogenase | P07014 |
| NG39 | dehydrogenase | P0DOV5 |
| NG40 | dehydrogenase | G3XD94 |
| NG41 | dehydrogenase | O69056 |
| NG42 | dehydrogenase | Q8X6C4 |
| NG43 | dehydrogenase | Q9ZGH1 |
| NG44 | methyltransferase | A8C927 |
| NG45 | methyltransferase | P9WJ63 |
| NG46 | methyltransferase | D5FKJ3 |
| NG47 | methyltransferase | P18644 |
| NG48 | methyltransferase | Q9F5K5 |
| NG49 | methyltransferase | Q9AJU1 |
| NG50 | methyltransferase | O87131 |
| NG51 | methyltransferase | A0A6B9HEI0 |
| NG52 | methyltransferase | P0C6Q8 |
| NG53 | methyltransferase | Q70KH3 |
| NG54 | methyltransferase | P36979 |
| NG55 | methyltransferase | Q9RMN9 |
| NG56 | methyltransferase | A0A0D4BS77 |
| NG57 | methyltransferase | A0A0F6P9C0 |
| NG58 | methyltransferase | P22610 |
| NG59 | methyltransferase | O54571 |
| NG60 | methyltransferase | O86262 |
| NG61 | methyltransferase | O31073 |
| NG62 | methyltransferase | Q51701 |
| NG63 | methyltransferase | A0A0H2ZF87 |
| NG64 | monooxygenase | P80645 |
| NG65 | monooxygenase | O87082 |
| NG66 | monooxygenase | Q59971 |
| NG67 | monooxygenase | Q9HTF3 |
| NG68 | monooxygenase | B0FXI0 |
| NG69 | monooxygenase | A0A0K2JL70 |
| NG70 | monooxygenase | P11295 |
| NG71 | monooxygenase | P9WKF7 |
| NG72 | monooxygenase | Q59723 |
| NG73 | monooxygenase | A4IU28 |
| NG74 | monooxygenase | A3KIM2 |
| NG75 | monooxygenase | A3KKC4 |
| NG76 | monooxygenase | Q838S1 |
| NG77 | monooxygenase | Q9I1C2 |
| NG78 | monooxygenase | A0QTU9 |
| NG79 | monooxygenase | P75898 |
| NG80 | monooxygenase | Q5YTV5 |
| NG81 | monooxygenase | P17055 |
| NG82 | monooxygenase | A0R4Q6 |
| NG83 | monooxygenase | Q08015 |
| NG84 | prenyltransferase | Q8GHB2 |
| NG85 | prenyltransferase | Q9L9F1 |
| NG86 | prenyltransferase | C4PWA1 |
| NG87 | prenyltransferase | A8M6W6 |
| NG88 | prenyltransferase | F9UT67 |
| NG89 | prenyltransferase | P0AG03 |
| NG90 | prenyltransferase | Q2L6E3 |
| NG91 | prenyltransferase | P69772 |
| NG92 | prenyltransferase | P69774 |
| NG93 | prenyltransferase | Q47RM6 |
| NG94 | prenyltransferase | D4G0R4 |
| NG95 | prenyltransferase | P0DV09 |
| NG96 | prenyltransferase | P33690 |

Table S3 Number of positive and negative sequences are balanced across functional categories.

| functional category | positive | negative |
| --- | --- | --- |
| AMP-binding enzyme | 76 | 52 |
| amidotransferase | 44 | 56 |
| dehydrogenase | 174 | 178 |
| methyltransferase | 143 | 110 |
| monooxygenase | 123 | 142 |
| prenyltransferase | 33 | 40 |
| sum | 593 | 578 |

Table S4 Detailed information of additional phenazine-interacting proteins in Test (II).

| label | gene | function | Uniprot | predidcted label |
| --- | --- | --- | --- | --- |
| positive | nalD | phenazine-binding repressor | Q9HY46 | positive |
| positive | mexR | phenazine-binding efflux pump regulator | A0A0C7ASE7 | positive |
| positive | ehpR | putative phenazine chaperon | Q8GPH6 | positive |
| positive | podA | pyocyanin demethylase | K0V2D8 | positive |
| positive | phdA | phenazine decarboxylase | A0A0N9Y7U2 | negative |
| positive | bphA1f | phenazine dioxygenase | A0ABV6CWE9 | positive |
| positive | pcaA1 | phenazine dioxygenase | A0A9J9LD27 | positive |
| positive | pcnH | phenazine amidase | A0A1T5GXF3 | negative |
| positive | pcnD | phenazine dioxygenase | A0A1T5GX78 | positive |
| positive | pzcH | phenazine amidase | UPI001A0572E3 | positive |
| negative | SFSIP | Ferric siderophore reductase | Q080S8 | positive |
| negative | pptP | 4'-phosphopantetheinyl transferase PptT | O33336 | negative |
| negative | norC | Nitric oxide reductase subunit C | Q59646 | negative |
| negative | ackA | Acetate kinase | P0A6A3 | negative |
| negative | DyP | Dye-decolorizing peroxidase | I6Y4U9 | negative |
| negative | rhlB | ATP-dependent RNA helicase RhlB | Q8P4D4 | negative |
| negative | xylA | Xylose isomerase | P24300 | negative |
| negative | pflB | Formate acetyltransferase | P09373 | negative |
| negative | lipA | Lipoyl synthase | P9WK91 | negative |
| negative | merA | Mercury(II) reductase | P00392 | negative |

Table S5 Primers used in this study.

| Name | 5’ - 3’ sequence | usage |
| --- | --- | --- |
| 3C-LP1 (pCoofy18_F) | GGGCCCCTGGAACAGAACTTCCAG | protein expression |
| ccdB-LP2 (pCoofy18_R) | CGCCATTAACCTGATGTTCTGGGG | protein expression |
| PTC_pCoofy18_F | AAGTTCTGTTCCAGGGGCCCATGTCAAACGATACCGGTTTCACTCATCTT | protein expression |
| PTC_pCoofy18_R | CCCCAGAACATCAGGTTAATGGCGCTATTATTTATTCAGCGATTGCGTGTTGCGG | protein expression |
| pBBR1MCS-2_F | CAATTCGCCCTATAGTGAGTCGTATTACGCG | complementation |
| pBBR1MCS-2_R | CAGCTTTTGTTCCCTTTAGTGAGGGTTAATTGCGC | complementation |
| PTC_pBBR1MCS-2_F | ACTAAAGGGAACAAAAGCTGATGTCAAACGATACCGGTTT | complementation |
| PTC_pBBR1MCS-2_R | ACTCACTATAGGGCGAATTGTTATTTATTCAGCGATTGCGT | complementation |
| kanR_PTC_F | ATCCTGCCTGTGGCATGGTGACAGCTCAACTTAAACAGGGAGCGGCCATGGTGTAGGCTGGAGCTGCTTC | deletion |
| kanR_PTC_R | GCAGTCACGCAACAAGTAATGGGATTCATTCTGTTCTCTCAACCGCTTTACATATGAATATCCTCCTTAG | deletion |
| PTC_upstrean_F | CTGACTGAATGCCTGCAGAACG | deletion |
| PTC_downstream_R | GGCCGCTCCCTGTTTAAGTTGA | deletion |
| PTC_upstream_F | AGCGGTTGAGAGAACAGAATGAATCC | deletion |
| PTC_downstream_R | CCTTGCTGCCCTGGAAGAAATG | deletion |

Dataset S1. Detailed information of all phenazine producer genomes (separate file).

Dataset S2. Amino acid sequences for all training data (separate file).

Dataset S3. Scores of each protein in Pantoea agglomerans genome (separate file).

Dataset S4. Detailed information of all homologs of PTC (separate file).
